# Discovering a less-is-more effect to select transcription factor binding sites informative for motif inference

**DOI:** 10.1101/2020.11.29.402941

**Authors:** Jinrui Xu, Jiahao Gao, Mark Gerstein

## Abstract

Many statistical methods have been developed to infer the binding motifs of a transcription factor (TF) from a subset of its numerous binding regions in the genome. We refer to such regions, e.g. detected by ChIP-seq, as binding sites. The sites with strong binding signals are selected for motif inference. However, binding signals do not necessarily indicate the existence of target motifs. Moreover, even strong binding signals can be spurious due to experimental artifacts. Here, we observe that such uninformative sites without target motifs tend to be “crowded” -- i.e. have many other TF binding sites present nearby. In addition, we find that even if a crowded site contains recognizable target motifs, it can still be uninformative for motif inference due to the presence of interfering motifs from other TFs. We propose using less crowded and shorter binding sites in motif interference and develop specific recommendations for carrying this out. We find our recommendations substantially improve the resulting motifs in various contexts by 30%-70%, implying a “less-is-more” effect.

## INTRODUCTION

Transcription factors (TFs) regulate gene expression by recognizing specific DNA sequences in the genome [1]. The targeted sequences of a TF share certain similarities, and thus can be summarized as a DNA motif. Identifying these motifs in the genome is the key to decipher gene regulation mechanisms and their evolution [2–5]. Although several *in vitro* techniques have been developed to determine TF binding motifs (TFBMs) [6, 7], many motifs are being inferred from *in vivo* TF binding sites identified by experiments such as chromatin immunoprecipitation followed by sequencing (ChIP-seq) [8, 9]. These inferred motifs account for a large fraction of known motifs in the major motif databases [10–12]. In addition, many widely used databases exclusively collect motifs inferred from the TF binding sites [13–17]. These many inferred motifs have become an important cornerstone of the field.

Operationally, motif inference aims to identify the statistically over-presented motifs from a set of TF binding sites. Its success depends on the designs of statistical models in use and the signal-to-noise ratio of the binding sites. Numerous statistical methods have been developed and tested during the past several decades, and developing new methods still receives much attention [16, 18–24]. To improve the signals in binding sites, auxiliary information such as GC content were used to reduce false-discovered sites [25–28]. However, the auxiliary information is difficult to be implemented in practice, and likely causes detection biases in motif discovery. Since ChIP-seq was invented, it has been the gold standard for detecting TF binding sites due to its higher accuracy and better resolution compared to its predecessors [29, 30]. Therefore, in the current common practice, the binding sites with strong binding signals determined by ChIP-seq are directly used for motif inference.

We argue that considering binding integrity alone may be insufficient for motif inference. First, TF binding may not be mediated by motifs, but instead by other factors such as enhancers, mediator proteins, or permissive DNA sequences [31–33]. Second, the sites with strong binding signals may also contain motifs of many other TFs (e.g. the motifs of co-binding TFs) [33–36]. Such motifs may interfere with the target motif inference [36]. These sites crowded with TFs tend to be from the high-occupancy target (HOT) regions which are widely observed in the genome [37–39]. In addition, even with improved accuracy, ChIP-seq may still generate many false positive sites [31, 32]. Such sites are usually a result of systematic biases in ChIP-seq rather than random technical noise, and thus can appear with strong artificial binding signals [39, 40]. Taken together, using strong ChIP-seq signals may not identify the most informative binding sites for motif inference.

A typical TF binding site from ChIP-seq is a 200 base-pair (bp) genomic region with a binding signal summit. Ideally, the target motif is close to the signal summit, however in practice, the actual distance depends on the summit accuracy and TF binding mode, and thus 200 bp or even longer genomic regions around the summits are often used for motif inference [41–43]. These lengths are much longer than a typical eukaryotic motif [44, 45], and thus likely include interfering motifs. Taken together, the optimal binding site length depends simultaneously on binding summit accuracy, TF binding modes, and interfering motif distribution in the site. Note that the interfering motifs we refer to are the binding motifs of other TFs rather than random motifs. The influence of such random motifs has been investigated both theoretically and empirically by simulating random sequences as pseudo-binding sites [46, 47]. Motif inference from the simulated sequences suggests that using shorter sites results fewer random motifs detected as false positive [47]. The signal of random motifs is usually weak, and thus can interfere mainly with weak biological motifs [46].

In sum, the widely used approach for binding site selection neglects two important issues: (1) the ambiguous association between binding signals and occurrences of target motifs and (2) the interference of other motifs with the target motifs. These two drawbacks probably cannot be fixed by improving ChIP-seq for TF binding prediction or using auxiliary information. We developed a simple computational approach to identify and filter out the noninformative binding sites that are susceptible to the two issues. In addition, we tuned the sites to find optimal lengths that provide the best tradeoffs between excluding interfering motifs and retaining target motifs. Finally, with this combined approach, we inferred accurate motifs for 1,207 TFs from their binding sites generated by the Encyclopedia of DNA Elements (ENCODE) [48] and model organism encyclopedia of regulatory networks (modERN) [49, 50] consortia.

## RESULTS

### Informative binding sites for motif inference

The binding sites of a TF detected by ChIP-seq can be classified into five categories (Fig. 1): (1) TF binding is mediated by its target motifs, and there are few motifs of other TFs in the binding site; (2) TF binding is also motif mediated, but many other TFs with their motifs concur within the site; (3) TF binding is mediated by a permissive DNA sequence rather than by the target motif, and the permissive sequence also recruits many other TFs; (4) the predicted TF binding site is spurious, and thus has no target motifs; and (5) TF binding is mediated by other TFs or enhancers, and thus there is no target motif in the site. Intuitively, the site in category 1 is more informative for motif inference than the others. However, using a binding signal from ChIP-seq may not necessarily enrich for sites from category 1.

**Figure 1.**
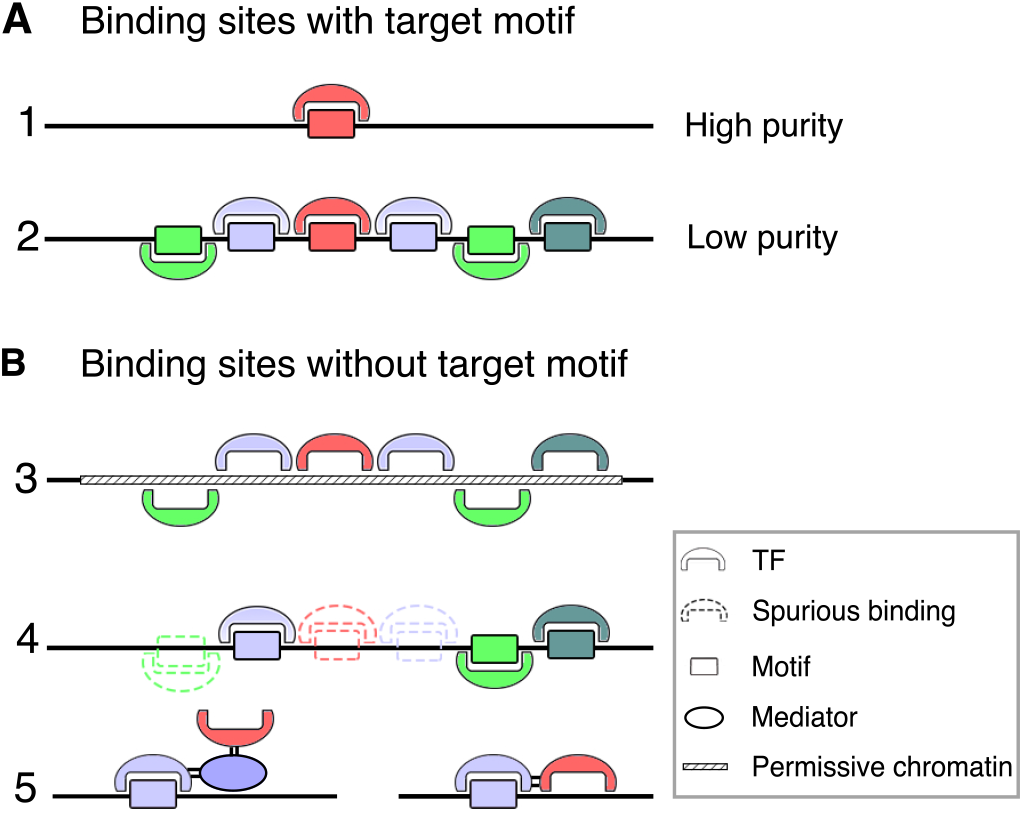
Classification of the binding sites of a target TF (red). The binding of the target TF can be mediated by its canonical motif which is denoted as a red rectangle (A). The binding site of the target TF may have few other TFs (A1) or many other TFs nearby (A2). The target TF binding can be mediated by a permissive genomic region, and due to its promiscuity, it also recruits many other TFs (B3). The target TF binding observed by ChIP-seq can be spurious due to systematic biases associated with a genomic region, and thus many other TFs also tend to have spurious sites observed in the region (B4). The target TF binding can be mediated by other factors such as mediator proteins and other TF factors instead of target motifs (B5).

Compared with category 1, the sites in categories 2-5 have more companion TFs. The TF crowdedness in categories 2, 3, and 5 is obvious by definition. For category 4, spurious sites are usually generated due to the properties of genomic regions; for instance, highly transcribed genomic regions contain many spurious sites [39, 40]. Therefore, spurious sites of different TFs tend to co-occur within such genomic regions, rendering artificial crowdedness. We postulated that filtering out sites with high TF crowdedness may improve motif inference by simultaneously enriching for target motifs and reducing interfering motifs. In addition, using short binding sites may further reduce interfering motifs in particular. We tested these two strategies individually and collectively for motif inference.

### Developing a crowding score from ChIP-seq data

Taking fruit fly, *Drosophila melanogaster*, as an example, we collected ChIP-seq data of the 452 unique fly TFs generated in the ENCODE portal. A binding site is defined as the 200 bp region centered on the binding signal summit determined by SPP software [51] in the standard ENCODE ChIP-seq pipeline. For each TF, we focused on its top 3,000 binding sites with strong binding signals. With the top binding sites of many TFs, the crowding score (C-score) of a site is simply the number of unique TFs with summits in the site. Similarly, we calculated C-scores for the binding sites of worms (*Caenorhabditis elegans*) and human cell lines (GM12878 and K562). A large C-score indicates that the site is likely from category 2-5, and thus is not informative for motif inference.

### Examining the effects of the crowding score on target motif enrichment

We focused on TFs with their target motifs determined by *in vitro* experiments such as SELEX and B1H in the Cis-bp database [52]. We took these motifs as the known motifs of the TFs, and calculated the motif enrichment of binding sites as the fraction of the sites with significant hits (*P* < 10^−4^) of target motifs by FIMO [53]. As a result, the sites with lower C-scores enriched for substantially more target motifs. Intriguingly, the C-score outperformed strong binding signal in motif enrichment for fly TFs (Fig. 2A). Particularly, the top 250 sites selected by C-score had 29.2% more target motifs than those selected by binding signal (Wilcoxon test, *P* < 8.4 × 10^−9^). For worm TFs, the C-score was also superior to the binding signal (Fig. 2B), and the improvement for the top 250 binding sites was 24.3% (*P* < 5.4 × 10^−4^). To test whether the improvements are due to the GC contents of the sites, we divided each site into 10bp bins, and then randomly shuffled the sequence within each bin. This broke any biological motifs within each site but remained the local GC contents. For these shuffled sites, the binding signals and C-scores have no difference in motif enrichment.

**Figure 2.**
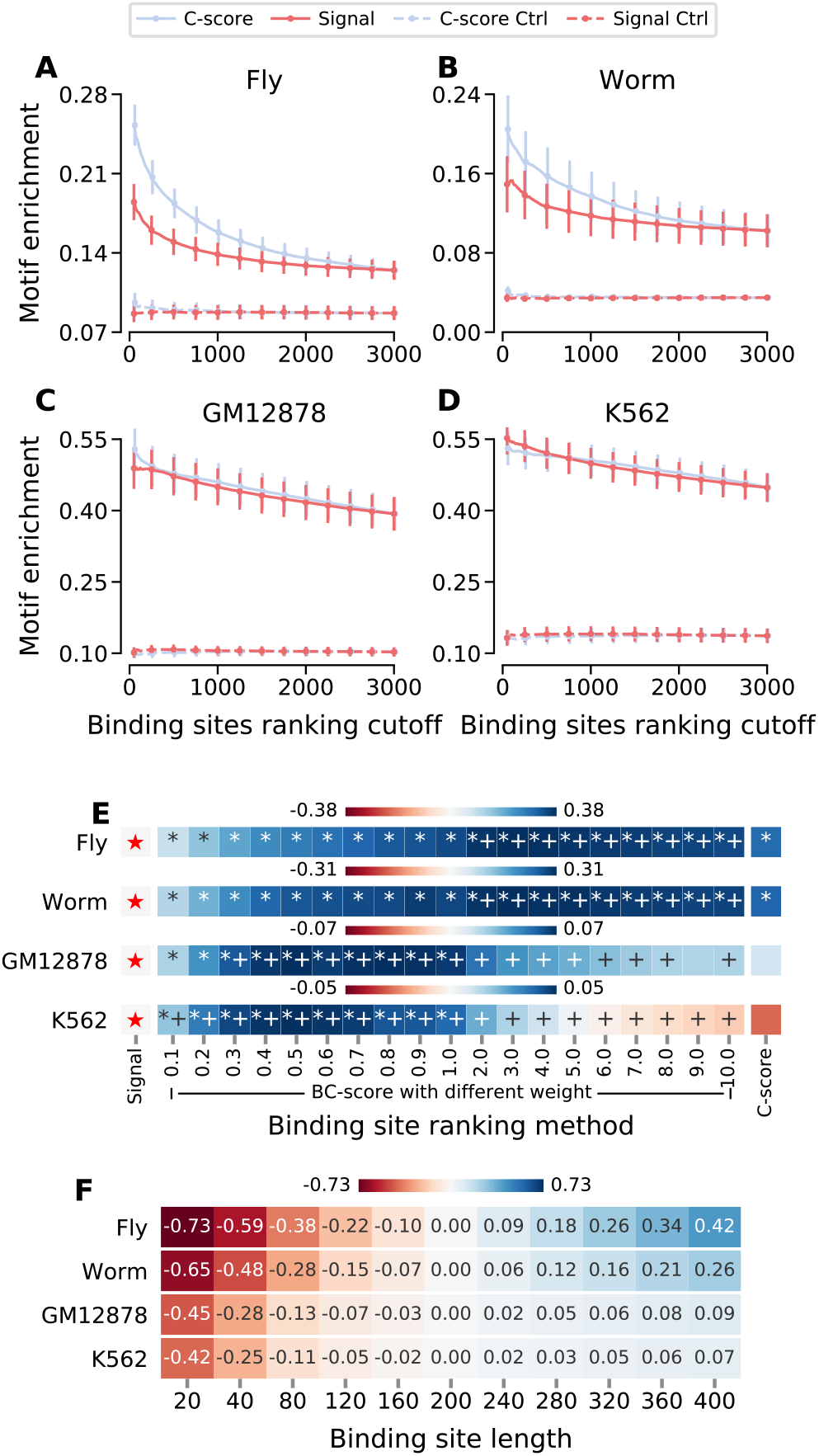
Motif enrichment in ranked binding sites and its sensitivity to binding site length. For a set of binding sites, the motif enrichment of a target TF is measured by the fraction of the sites containing the known (*in vitro*) motifs of the TF. This enrichment is calculated for the top binding sites ranked respectively by binding signal from SPP, crowding score (C-score) and binding-crowding score (BC-score). The results respectively for fly, worm, GM12878 and K562 are showed in the panels A to D, in which the dots and fitting lines indicate the motif enrichments averaged over the TFs. The whiskers indicate the standard errors. The dash lines indicate the enrichment in shuffled sites. In panel E, BC-scores with different weights on C-score are used to rank the binding sites of TFs. The motif enrichment in the top 250 binding sites is compared to that in the top 250 binding with strongest binding signals as a reference (denoted by a red star). The difference is measured by the fraction change between the averages in motif enrichment. Wilcoxon tests are used, and sample sizes are 196, 59, 62 and 99 for fly, worm, GM12878 and K562, respectively. ‘*’ indicates that the motif enrichment by the BC-score is significantly different from that by the binding signal. ‘+’ indicates that the motif enrichment by the BC-score is significantly different from that by the C-score alone. In panel F, for the top 250 binding sites with strongest binding signals, the binding site length is tuned around 200 bp, and the fractional change in average motif enrichment is calculated compared to that of 200 bp.

For the TFs in the human cell lines, the C-score and binding signal performed similarly in motif enrichment (Fig. 2C&D), suggesting that the human TFs have less sites from categories 3-5 compared to fly and worm TFs. This is consistent with our discovery that fly and worm TFs have more spurious sites (category 4) than human TFs, which is likely due to the difference in genome activity between fly/worm and human cell lines [54]. Although the C-score performed similar to the binding signal for human TFs in motif enrichment, it is expected to reduce interfering motifs substantially; thus, the C-score would likely improve motif inference. Moreover, combining the C-score and the binding signal may further improve target motif enrichment.

### Combining the crowding score and the binding signal of ChIP-seq

We first ranked the sites of a TF by their binding signals and C-scores, respectively, in descending and ascending order. For a binding site *i*, the ranks in binding signal and C-score are denoted respectively as *b*_*i*_ and *c*_*i*_. Therefore, the binding-crowding score (BC-score) of the site is *b*_*i*_ + *w* × *c*_*i*_, where weight (*w*) is a parameter to determine the contribution by the C-score. A *w* larger than one indicates a higher belief in C-score than binding signal. We used *w* ranging from 0.1 to 10 for the BC-score to rank binding sites. For the top 250 sites ranked by BC-scores, many different *w* can further improve target motif enrichments, compared to using binding signals and C-scores alone (Fig. 2E). As expected, the optimal *w* on C-score is around 3.0 for both fly and worm TFs, which is much larger than those for human TFs (Fig. 2E). Note that for human TFs, the BC-scores with multiple weights also significantly outperformed the binding signal in target motif enrichment.

### Examining the impact of binding site length on target motif enrichment

We postulate that truncating binding sites around summits may lose target motifs substantially. To test this, we focused on the top 250 binding sites with strong binding signals. Indeed, for fly and worm TFs (Fig. 2F), the target motif enrichment reduced rapidly with reducing the binding site length. However, for human TFs, the reduction rate was minimal (Fig. 2F). Taking the change from 200 bp to 120 bp as an example, this 40% shorter length reduced the motif enrichments by 22% and 15% respectively for the fly and worm TFs, whereas the resultant reductions were only 5%-7% for the human TFs, suggesting that using shorter binding sites may benefit the human samples more than the fly/worm samples in motif inference.

### An integrative framework to select informative binding sites for motif inference

To integrate binding site length and BC-score together for motif inference, we first used the BC-score (*b* + *w* × *c*) to identify the top 250 binding sites of a TF, and then truncated the selected sites around their summits to a length *l*. The weight *w* and length *l* are the only two parameters that need to be tuned for binding site selection. For comparison, we took the conventional binding site selection as the reference, i.e. the top 250 binding sites with strong ChIP-seq signals and site lengths equal to 200 bp centered on the summits. This number of binding sites used is reasonable in this study because it falls in the transition regions of the curves in Figure 2, and thus likely maximizes motif enrichments and sample sizes for high statistical power. From our selected sites and the conventional sites, we can infer binding motifs respectively and compare them to the motifs determined by *in vitro* experiments for evaluation.

To infer motifs from a set of binding sites, we used the recently developed and widely used software DREME [18]. For each TF, the top 5 inferred motifs were compared to the *in vitro* motifs using TOMTOM [55]. Using the conventional binding site selection, on average only 0.684 out of the five inferred motifs are significantly similar (*P* < 0.05) to the known target motifs of the fly TFs. Using the C-score alone, this number increased substantially by 31.3% to 0.898 (*P* < 3.1 × 10^−2^, Fig. 3A). In addition, with a wide range of weights on C-score, the BC-scores further improved the motif inference; the best improvement was 47.8% with the weight equal to 5.0 on C-score (*P* < 2.0 × 10^−3^, Fig. 3A). Finally, simultaneously optimizing both binding site lengths and BC-scores enhanced the improvement to 71.6% (*P* < 1.3 × 10^−4^, Fig. 3A).

**Figure 3.**
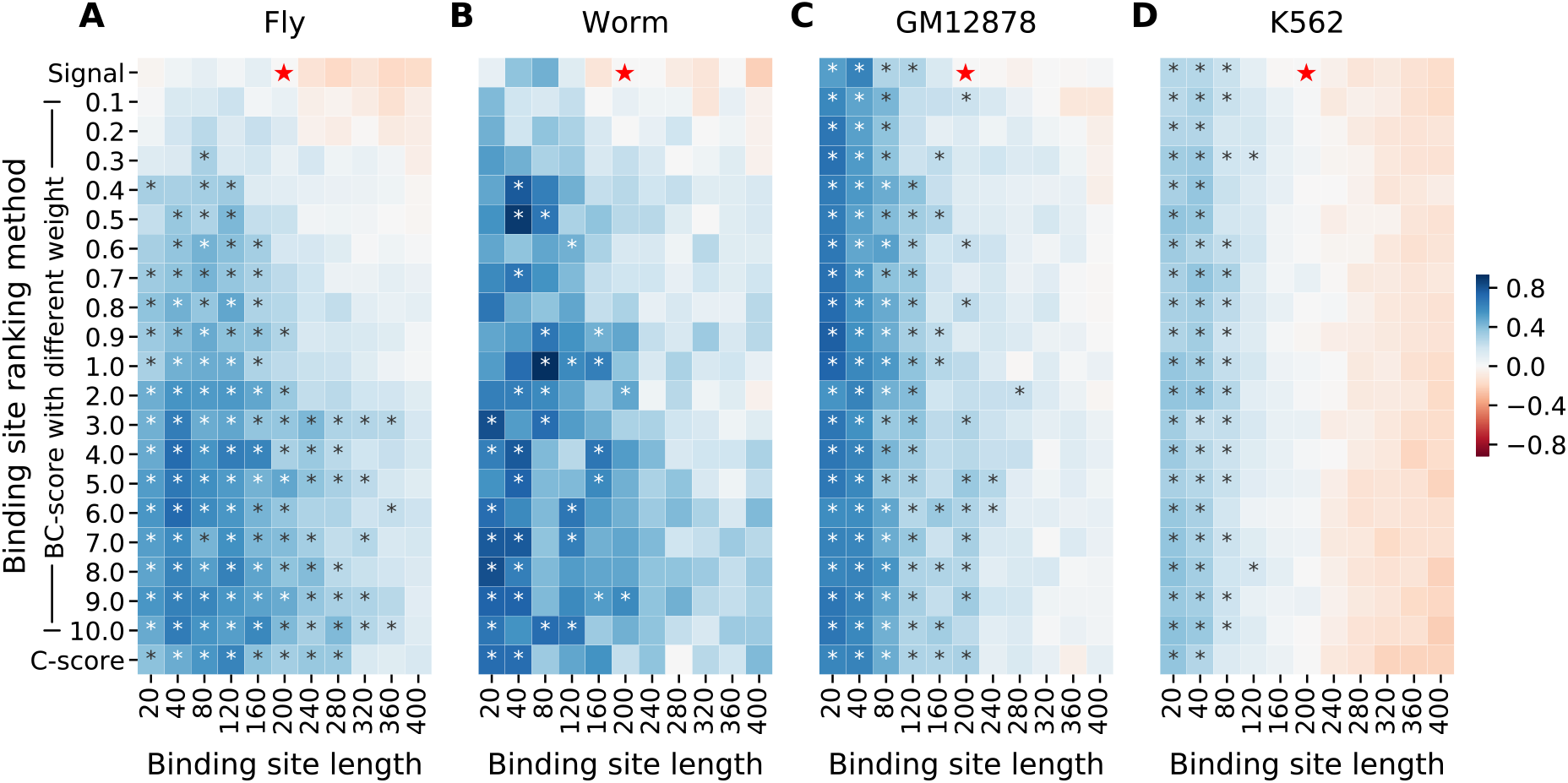
Fractional differences in motif inference between the conventional binding site selection and our selection. The conventional selection uses the top 250 binding sites with strongest ChIP-seq binding signals for motif inference. The site lengths are equal to 200 bp. Our selection uses C-scores or BC-scores to select binding sites, and the site lengths are tuned between 20bp to 400bp. From the selected sites of each TF, DREME is used to infer five motifs. The number of the five motifs that are significantly similar to the *in vitro* motifs indicates the accuracy of the motif inference. For the fly TFs (A), the red star indicates the conventional selection, and each of the other grids indicates the selection method using certain parameters. The color of a grid indicates the fractional difference in the average accuracy between the corresponding selection method and the conventional selection. ‘*’ indicates *p* < 0.05 from Wilcoxon tests. The sample sizes are 196, 59, 62 and 99 for fly (A), worm (B), GM12878 (C), and K562 (D), respectively.

For the worm TFs, C-score alone also substantially improved motif inference by 22.2%; however, this improvement was not statistically significant (*P* = 0.23, Fig. 3B), presumably due to the much smaller sample size of worm (59) than that of fly (196). Nevertheless, BC-scores with various weights on C-score both substantially and significantly improved the motif inference. The optimal weight of the C-score in the BC-score was 9.0, resulting in a 59.3% improvement in motif inference (*P* < 4.5 × 10^−2^, Fig. 3B). Furthermore, optimizing the site length increased the improvement to 92.6% (*P* < 5.2 × 10^−3^, Fig. 3B). Similarly, for the human TFs, BC-scores with different weights substantially improved motif inference. With the weights equal to 5.0 and 3.0, the best improvements were 34.0% and 6.59% for GM12878 and K562, respectively. With optimal site lengths, these improvements increased to 75.5% and 37.9%, respectively.

As expected, truncating the top binding sites with strong binding signals benefited the motif inference more for human TFs (Fig. 3C&D) than for worm and fly TFs (Fig. 3A&B) because the improvement of the human motif inference is mainly associated with the binding site length. For human TFs, binding site length was the main factor for improving the motif inference. As for fly and worm TFs, both shorter binding sites and larger weights on C-score improved the motif inference substantially (Fig. 3A&B). However, for all the species, the best performance requires tuning both factors. By their definitions, both low C-scores and short binding sites reduce interfering motifs. Due to this partial redundancy, we observed that a certain improvement can be achieved by either tuning the weight on C-score or the binding site length (Fig. 3A&B), indicating multiple sets of optimal parameters available for motif inference. The **mo**tif inference using our **bi**nding site selection approach is referred to as MOBI.

Estimating the C-scores of a sample such as GM12878 or K562 requires a large number of TFs with binding sites detected by ChIP-seq. However, many samples do not have sufficient ChIP-seq data. Therefore, we tested whether pooling the ChIP-seq data of multiple samples can generate informative C-scores for another sample. To this end, we calculated C-scores from the ChIP-seq data of all the cell lines (521 TFs) except GM12878 (136 TFs) and K562 (336 TFs), and then used these C-scores to infer motifs for GM12878 and K562, respectively. For GM12878, with many weights, these transplanted C-scores improved the motif inference substantially, compared to the conventional inference (Fig. S1). However, for K562, an insignificant improvement was observed only when the C-scores had a small weight (Fig. S1). Taken together, these results indicate that transplanted C-scores are not robust for motif inference.

### Cross validating the MOBI approach for motif inference

Taking the fly TFs as an example, we randomly divided the TFs with known motifs into five groups. Four of them were pooled as a training dataset, and the remaining group served as a testing dataset. For the training TFs, we first identified the optimal values of *w* and *l* for motif inference using DREME, and then validated the performance of these two values on the testing TFs. We repeated this procedure multiple times to have a robust estimation of the MOBI approach. From each of the testing sets, we also conducted the conventional motif inference, i.e. using top sites with strong binding signals and 200bp lengths. On the testing datasets, the MOBI approach with DREME improved motif inference by 60.7% on average, compared to the conventional motif inference (Fig. 4A). For worm, GM12878, and K562, the average improvements are 63.6%, 66.2%, and 26.4% respectively (Fig. 4A).

**Figure 4.**
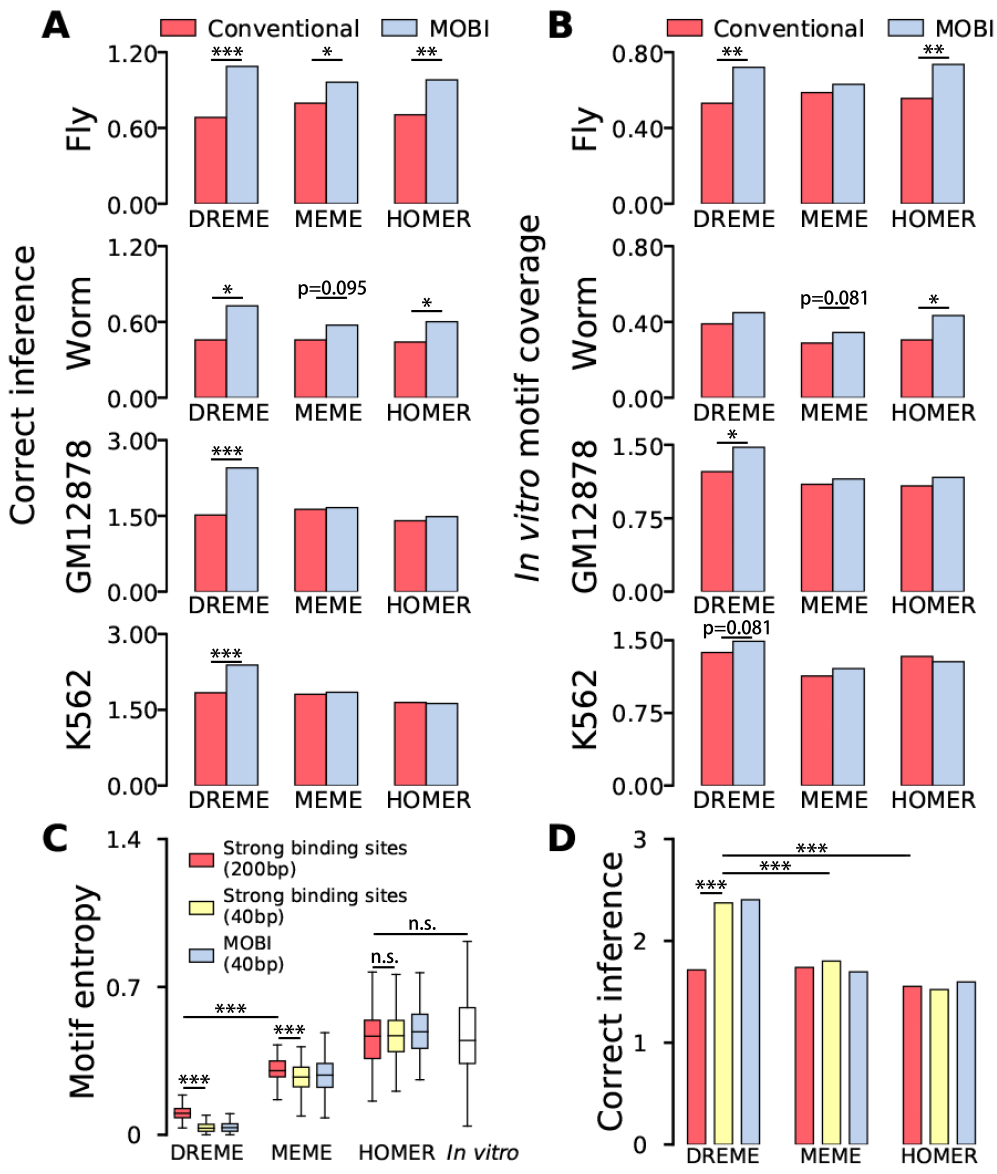
Validating the performance of MOBI binding site selection for motif inference. Two measures are used to evaluate the motif inference for a TF. The first one is the number of the inferred motifs that are significantly (*p* < 0.05) similar to the *in vitro* motifs, referred to as correct inference (A). The other is the number of the *in vitro* motifs that are significantly similar to the inferred motifs, referred to as coverage (B). The available TFs from each sample (e.g. GM12878 cell line) are divided evenly into a training set and a testing set. The MOBI approach first identifies its optimal C-score weight and the site length, which result in the best correct inference for the training set. With these optimal parameters, the MOBI approach then infers the motifs for the TFs in the testing set. The resultant correct inference and coverage are compared to those of the conventional selection for the testing set. The conventional selection uses only the strong binding sites of the testing TFs, and the binding site length is 200 bp. With bootstrapping, this validation process was repeated 4,000 times for significance, i.e. the frequency that the conventional selection outperforms MOBI. The entropies of the inferred and *in vitro* motifs are calculated and compared, and high entropies indicate diverse DNA sequences in the motifs (C). The correctly inferred motifs associated with the entropies are shown in panel D. ‘*’, ‘**’, and ‘***’ indicate the p-values lower than 0.05, 0.01, and 0.001.

Compared to conventional binding site selection, our MOBI selection resulted in more motifs inferred correctly averaged across the TFs. Moreover, the MOBI selection also resulted in a larger fraction of the TFs with improved motif inference (Fig. S2). In addition, for each TF we examined the number of known target motifs that are significantly similar to the five inferred motifs, herein referred to as coverage. For this measure, our MOBI approach again outperformed the conventional selection substantially. The average improvements on the testing tests were 36.2% and 22.8% for the fly and worm TFs, respectively (Fig. 4B). For TFs in GM12878 and K562, the improvements were 26.5% and 5.99% (Fig. 4B).

### Interaction between MOBI and inference methods

In addition to DREME, we also used two popular methods, i.e. HOMER [41] and MEME [19], to repeat the cross-validations. Consistently among the three tools, our MOBI approach for binding site selection substantially outperformed the conventional selection in motif inference for the fly and worm samples (Fig. 4&S2). However, for the human cell lines, only with DREME, the MOBI approach outperformed the conventional selection, and this improved motif inference with DREME is also substantially better than those of the other tools using either the MOBI or conventional selection. Taken together, these results suggest that particularly for the human cell lines, HOMER and MEME do not benefit substantially from our MOBI approach. For the cell lines, the MOBI approach only reduces interfering motifs but does not substantially increase target motif enrichment in binding sites (Fig. 2C, D&E). Therefore, we postulate that different from DREME, MEME and HOMER tend to prefer diverse sequences for a motif regardless of the binding-site complexity, resulting in low-accuracy motifs.

To test this postulation, we calculated and compared the entropies of motifs. A high entropy indicates that the binding sequences of a motif are very diverse. Inferred from the top binding sites with strong binding signals and 200bp lengths, the DREME motifs have substantially lower entropies than those MEME or HOMER motifs (Fig. 4C). With the site lengths reduced to 40bp, the entropies of DREME motifs further decreased significantly, whereas those of MEME and HOMER motifs remain or even increase (Fig. 4C). Although the MEME and HOMER motifs have similar entropies to the *in vitro* motifs, the *in vitro* motifs are less similar to the MEME and HOMER motifs than to the DREME motifs (Fig. 4D). These results suggest that the high entropies of the MEME and HOMER motifs are likely due to including spurious binding sequences. The motifs inferred by the MOBI approach (i.e. *l* = 40bp and *w* = 0.9) using the three software have the same entropy patterns as the ones from the short sites. Note that low entropy does not artificially result in high motif similarity because the DREME motifs inferred from the conventional sites (200bp) also have substantially lower entropies than the MEME motifs, but these two groups of motifs have the same similarities to the *in vitro* motifs.

### Interaction between motif inference and sample specificity

With the conventional binding site selection, K562 seemingly has better motif inference than GM12878 (Fig. 4). These two human cell lines provide an opportunity to investigate the influence of genomic properties on motif inference. To this end, we focused on DREME and the common TFs that have *in vitro* motifs and ChIP-seq data in both GM12878 and K562. The top sites with strong binding signals from K562 enriched for more target motifs than those from GM12878 (*P* = 0.06, Fig. 5A), which likely resulted in the better motif inference observed for K562 using the conventional selection (*P* = 0.04, Fig. 5B). The higher motif enrichment is indeed significantly associated with the better motif inference (*P* = 0.05, Fisher exact test). Compared to GM12878, this higher motif enrichment in K562 may be facilitated by its more accessible genome to TFs [54]. Because the K562 binding sites with strong signals are already highly enriched with target motifs, using our MOBI approach results in a relatively small improvement for K562 (Fig. 5B).

**Figure 5.**
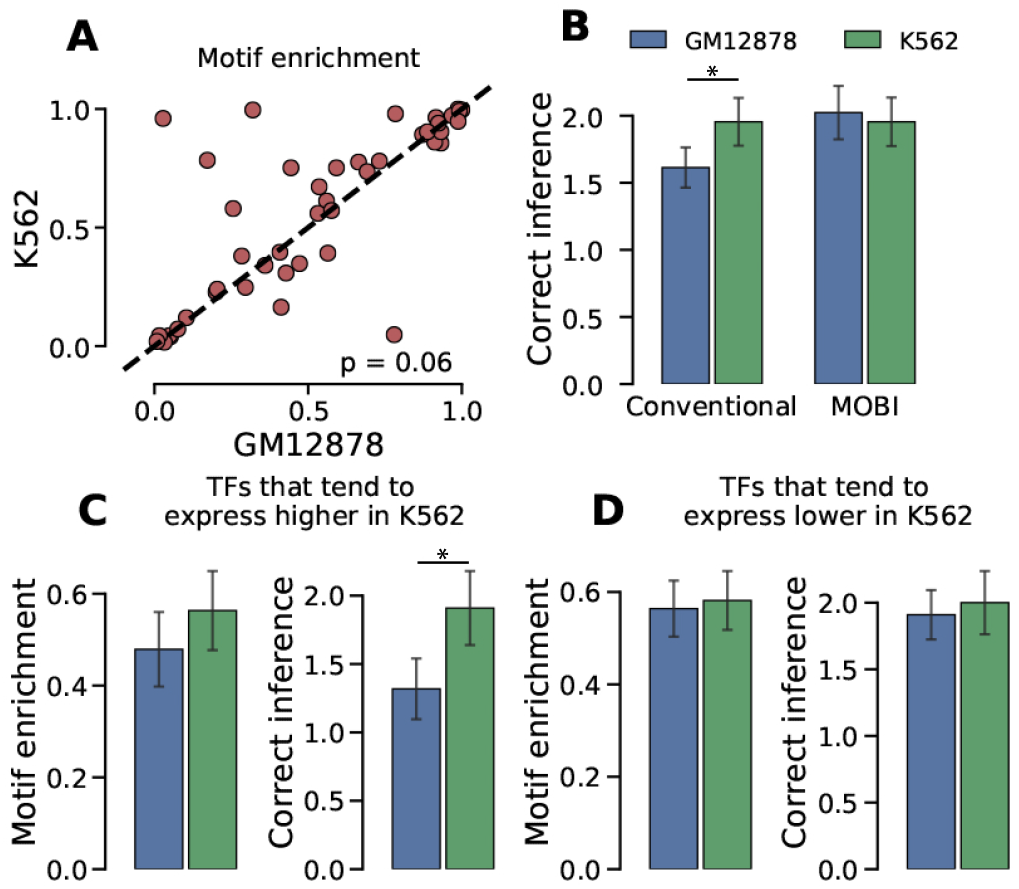
Comparing GM12878 and K562 for target motif enrichment and motif inference. For the common TFs between GM12878 and K562, the top 250 binding sites with strongest binding signals detected in K562 enrich for more target motifs than those detected in GM12878 (A). The conventional selection and the MOBI approach are used to infer the motifs of the common TFs (B). Wilcoxon tests are used with sample sizes equal to 44. The common TFs are then divided into two groups with equal numbers. One group contains the TFs that tend to have higher expression in K562 than in GM12878. With the conventional selection, the binding sites of such TFs detected from K562 enrich for more target motifs than those detected from GM12878 and also have better motif inference (C). For the rest TFs, their binding sites respectively from K562 and GM12878 are very similar in motif enrichment and correct inference (D). Wilcoxon tests are used, and the sample sizes are 22.

In addition to the genome accessibility, we investigated whether TF expression likely has detectable effect on motif inference. To this end, for each of the common TFs we calculated the ratio of its expressions in K562 and GM12878, and then ranked the TFs according to their ratios in descending order. We assumed that the top 50% TFs would tend to have lower expression in GM12878 than in K562. As a result, the strong binding sites of the top 50% TFs enriched fewer target motifs in GM12878 than in K562 (*P* = 0.10, Fig. 5C), and likely resulted in the less accurate motif inference (*P* = 0.03, Fig. 5C). Taking the bottom 50% TFs as a control, their target motif enrichment (*P* = 0.29) and motif inference (*P* = 0.32) have no significant difference between K562 and GM12878 (Fig. 5D). Probably because as a cancer cell line, K562 has high genome accessibility and most genes highly expressed, the top 50% and bottom 50% TFs in K562 have similar accuracy in motif inference (Fig. 5C&D).

### Inferring motifs for the TFs in the ENCODE portal

We demonstrated that with DREME, the motif inference using the MOBI approach outperformed the one using the binding sites with strong binding signals, referred to as the baseline. However, the baseline we implemented may have low accuracy, rendering the improvement not very useful. To test this, we compared the baseline to other databases of inferred motifs, i.e. Factorbook [56] and the one built by Kheradpour and Kellis [13]. These databases contain motifs inferred from the TF binding sites detected by the ENCODE consortium. For the common TFs with known motifs, our baseline performed better than Factorbook (*P* < 3.5 × 10^−4^; Fig. 6A) and similarly to the database by Kheradpour and Kellis (*P* = 0.18; Fig. 6B), which integrated multiple software for the motif inference. Taken together, these results indicate that our baseline is comparable to existing inferred motifs, and thus the motifs inferred by our MOBI approach are expected to be much more accurate. For each of the samples, i.e., fly, worm, GM12878, and K562 in ENCODE, we used its TFs with known motifs to identify the optimal MOBI parameters (i.e. site length and C-score weight), and then used these parameters to infer the motifs for all the TFs of the sample. As a result, we generated accurate motifs for the 1,207 TFs of human, worm, and fly.

**Figure 6.**
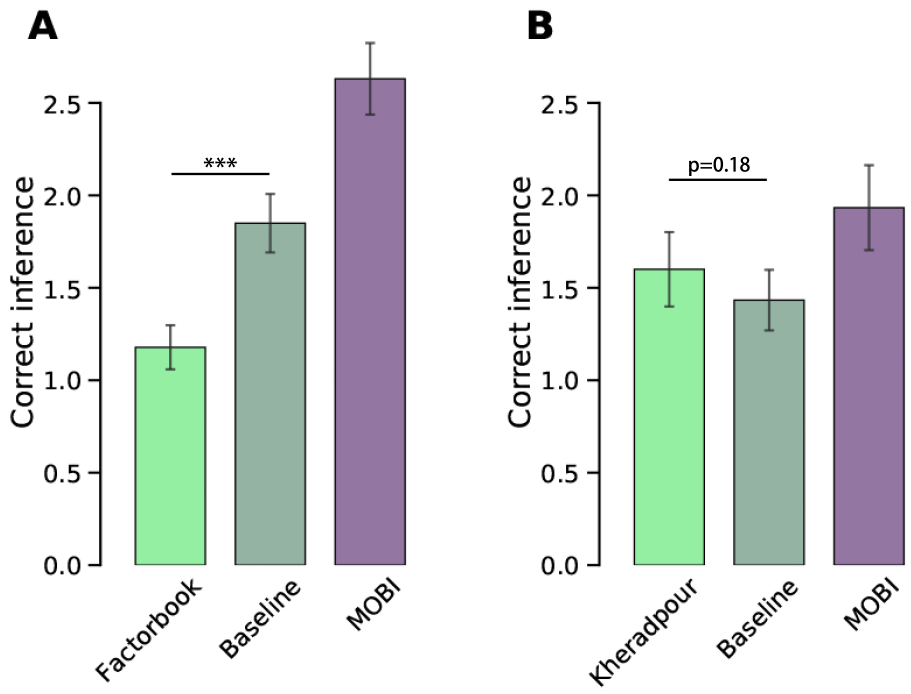
Comparing the motif inferences of our baseline to those of Factorbook (A) and Kheradpour & Kellis (B). Wilcoxon tests are used with sample size equal to 73 for factorbook and 30 for the Kheradpour & Kellis dataset.

### Inconsistency between motif enrichment and conventional motif inference

DNA input is the predominantly used control experiment in ChIP-seq. However, complex samples such as fly and worm whole organisms tend to have many spurious sites due to the nonspecific interactions between the antibodies and nonantigens, and thus another type of controls, mock IP, perform better for these complex samples [54]. With mock IP controls, we predicted the binding sites for a subset of the fly and worm TFs [54]. Indeed, the binding sites by mock IP controls enriched for significantly more target motifs than those by DNA input controls (Fig. S3). Surprisingly, for worm TFs, the higher enrichment of target motifs in the mock IP sites resulted in less accurate motif inference (Fig. 7A). This inconsistency may be because the mock IP sites also enriched for more interfering motifs. To this end, we used C-scores, which by design filter out interfering motifs, to identify the informative sites. With C-scores, the selected sites predicted by mock IP had higher enrichment of target motifs and more accurate motif inference than those predicted by DNA input (Fig. 7B). For the binding sites predicted by mock IP controls, using MOBI approach also substantially improved motif inference for both fly and worm TFs (Fig. 7C&D).

**Figure 7.**
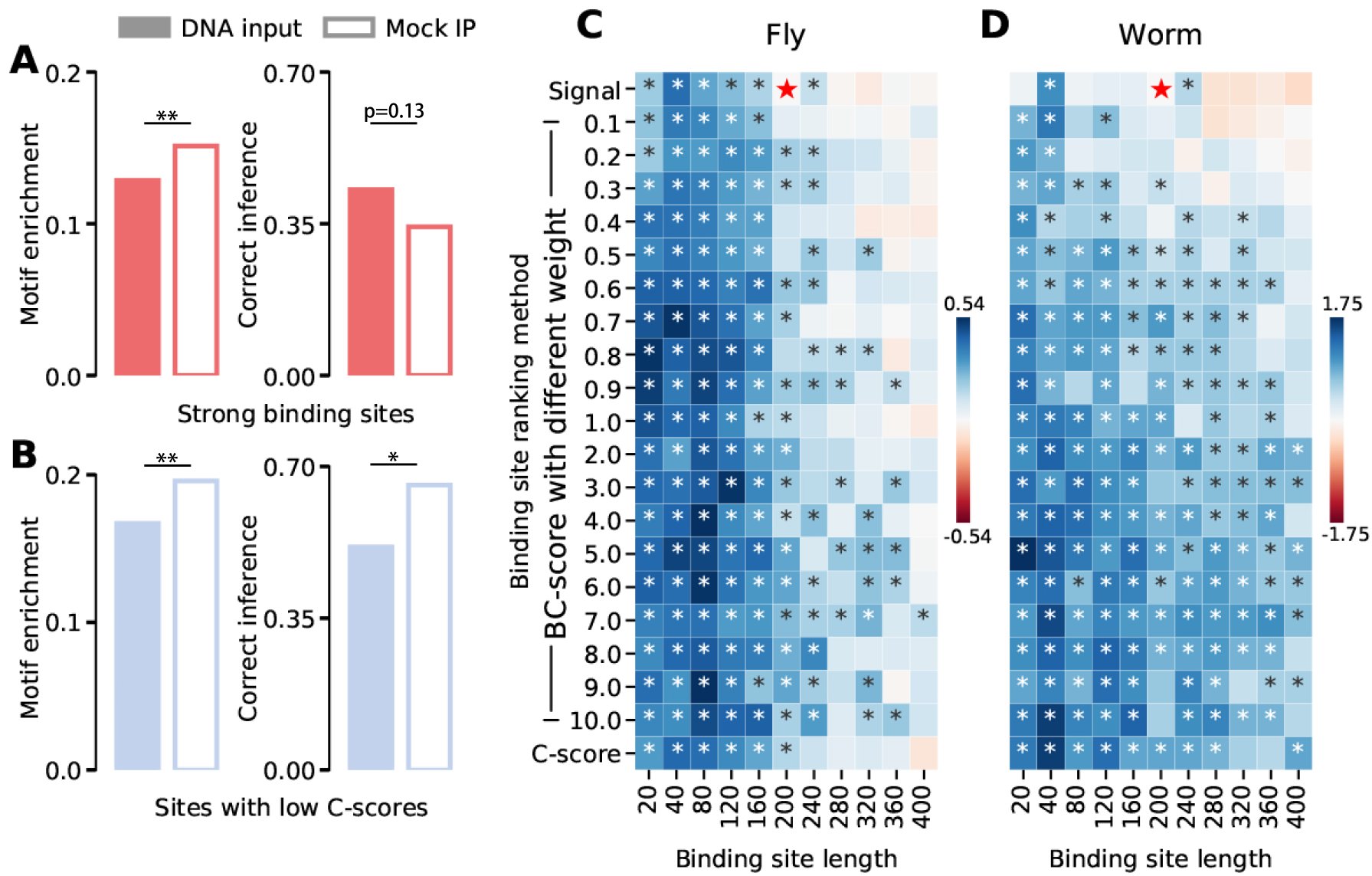
Interaction between binding site accuracy and motif inference. For the complex samples such as worm whole organism, the TF binding sites from ChIP-seq using mock IP controls have significantly higher target motif enrichment than those from ChIP-seq using DNA input controls, but the high target motif enrichment does not result in better motif inference, presumably due to interfering motifs (A). In line with this, when C-scores are used to select binding sites, the target motif enrichment and motif inference are both improved consistently (B). ‘*’ indicates statistical significance (*p* < 0.05) from Wilcoxon tests. For the ChIP-seq with mock IP controls, our MOBI approach still substantially improves motif inference, compared to the conventional binding site selection, for both fly TFs (C) and worm TFs (D). The conventional selections are denoted by red stars. ‘*’ indicates statistical significance (p < 0.05) from Wilcoxon tests. The sample sizes for fly and worms are 140 and 37 TFs.

## DISCUSSION

Informative binding sites and statistical methods are two pivotal factors in motif inference. Numerous statistical methods have been developed, whereas to our knowledge, the only widely used binding site selection for motif inference focuses on the sites with strong binding signals. The success of this selection relies on two assumptions: (1) the binding signal is strongly correlated with the existence of the binding motifs used by the TFs and (2) interfering motifs are not prevalent in the sites with strong binding signals. For the first assumption, we demonstrated that sites with strong binding signals can have low enrichment of the target motifs for two reasons. First, a large fraction of TF binding with high affinity may be mediated by enhancers and mediators instead of target motifs. Second, many strong binding signals may be spurious due to systematic bias uncontrolled in ChIP-seq [39, 40, 54]. The latter case is particularly prevalent in sites detected from samples with highly transcribed genomes [54].

For the second assumption, we found that interfering motifs in the sites with strong binding signals are abundant enough to impair motif inference. This is in line with the prevalent observation that different TFs bind to the same genomic regions to work cooperatively [57]. In fact, there is motif inference software developed particularly for detecting such co-binding motifs [18]. The binding sequences of the co-binding TFs may have similarities, and thus the inferred motifs are mixtures of the binding sequences for different TFs. In addition, the co-binding sites likely have strong binding signals, and thus tend to be used for motif inference. For example, the TFs of interest may be stabilized by the co-binders interacting with the TFs or creating favorable chromatin states for the TFs. As a result, selecting binding sites with strong binding signals may enrich for interfering motifs. We developed the C-score to exclude such sites that contain many interfering motifs and/or no target motifs. Particularly for interfering motifs, we tuned binding site lengths to exclude interfering motifs without losing many target motifs.

In genomes, cis-regulatory elements such as promoters and enhancers often contain multiple binding sequences for the same TF [58, 59]. These sequences can create multiple opportunities to recruit the TF, which results in robust and thus strong TF binding [60]. This redundancy for robustness is widely observed in both prokaryotes and eukaryotes [61]. In line with this, we observed that for the top binding sites detected from the human cell lines, the sites with stronger ChIP-seq signals tend to contain more motifs (Fig. S4). In addition, the motifs inferred by MOBI approach likely include the preferred motifs by the TFs, if there are any, because the MOBI approach usually focuses on the short sequences around the summits of binding signals, and such regions presumably correspond to the DNA sequences preferred by the TFs.

We demonstrated that for the binding sites predicted from the fly and worm whole organism, the MOBI approach can enrich for target motifs and reduce interfering motifs, and thus the three popular inference tools, namely, DREME, MEME and HOMER, all benefit from using this approach. For the binding sites from human cell lines, the MOBI approach only reduce interfering motifs, and the three tools benefit quite differently from MOBI. This difference is likely due to the intrinsic algorithms of the tools. With more inference tools developed [20–22, 62, 63], our MOBI approach may further improve motif inference. Compared to ChIP-seq, the recently developed ChIP-exo and ChIP-nexus are technically difficult and costly [64] but have much higher resolution, resulting in short binding sites [65, 66]However, same as ChIP-seq, these new techniques probably still cannot filter out the sites mediated by factors other than binding motifs. With sufficient data generated by these techniques, the MOBI approach is expected to infer more accurate motifs.

We inferred motifs for 1,207 TFs in the ENCODE portal, resulting in the MOBI database. Due to using our binding site selection, these inferred motifs are more accurate than those of the other databases [13, 56]. Moreover, our inferred motifs cover not only unique human TFs (375) but also many unique TFs of worm (283) and fly (452). These uniformly inferred motifs facilitate evolutionary analyses across species. The success of our binding site selection depends partially on C-scores. We found that the C-score is moderately correlated with genomic accessibility calculated from DNase-seq. Therefore, we used genomic accessibility instead of the C-score for binding site selection (Supporting information, Fig. S5). Although this genomic accessibility does not outperform the C-score, the inference results indicate that, nonintuitively, the highly accessible genomic regions we observed tend to be uninformative for motif inference.

In sum, we demonstrated that binding sites with strong binding signals can be far from optimal for motif inference. To this end, we developed the MOBI approach to identify informative binding sites for motif inference. Compared to the conventional binding site selection, this approach substantially improves motif inference. In general, the improvement due to the novel binding site selection is at least comparable to the difference in motif inference using the three inference methods, indicating that binding site selection may warrant more attention. With our MOBI approach, we inferred more accurate motifs for 1,207 TFs from human, worm, and fly. These motifs from human and the model organism are a valuable resource for comparative analyses of TF binding and its evolution.

## MATERIALS AND METHODS

### ChIP-seq data from the ENCODE portal

All ChIP-seq data in use were downloaded from the ENCODE portal. For human (genome build hg19), we used the ChIP-seq data of 136, 366, and 521 TFs respectively from GM12878, K562, and all the other cell lines, generated by ENCODE [67]. For fly (genome build dm6) and worm (genome build ce11), we collected ChIP-seq data for 452 and 283 TFs, generated by the modERN and modENCODE projects [50]. In addition, we used 304 and 168 ChIP-seq data with mock IP controls, generated for the fly and worm samples [54]. If multiple ChIP-seq datasets were available for a single TF, we randomly selected one of them for analysis. We defined a binding site as a 200 bp genomic region centered on a binding signal summit determined by the SPP pipeline [51]. Only the top 3,000 binding sites ranked by the binding signal were kept for further analysis. For TFs with less than 3,000 binding sites, all binding sites were kept.

### Developing MOBI approach for binding site selection

For a binding site, we define its crowding score (C-score) as the number of binding summits of unique TFs in the site. Respectively for the samples we used, namely GM12878, K562, and worm/fly whole organism, we calculated C-scores using the binding sites of the TFs with ChIP-seq data from the sample. The binding sites of a TF are ranked respectively according to their binding signals from ChIP-seq (in descending order) and the C-scores (in ascending order). Ties in C-score ranking are resolved by binding signals. For a binding site *i*, its ranks in binding signal and C-score were *b*_*i*_ and *c*_*i*_. The binding-crowding score (BC-score) of the site is thus denoted as *b*_*i*_ + *w* × *c*_*i*_, where *w* is a weight on the C-score rank. The BC-score with various weights was used to select informative binding sites. These sites were then trimmed around their summits to have a certain length, *l*. *l* and *w* were tuned to identify the most informative binding sites for motif inference. This binding site selection approach is referred to as MOBI.

### *In vitro* TF binding motifs and motif enrichments

We collected fly, worm, and human TF binding motifs determined by *in vitro* experiments, including SELEX, PBM, and B1H, from the Cis-bp database [52]. For the motifs of a TF, we concatenated all the unique position-weighted matrices into a single file in MEME motif format. As a result, 196 of the fly TFs in use have such known motifs. For GM12878, K562, and worm, these numbers were 62, 99, and 59, respectively. FIMO (v4.11.4) [53] was then used to scan the genomes for significant hits of the known motifs, using --max-stored-scores=10000000 and a nominal *P* value < 10^−4^. For a set of binding sites, the fraction of binding sites with significant motif hits was used to indicate the motif enrichment. To test the effects of GC contents, we separated the whole genome into 10 bp bins and randomly shuffled the sequence within each bin, and then scanned the known motifs in the shuffled genome and calculated motif enrichments for the binding sites.

### Motif inference and evaluations

We ranked the selected binding sites of each TF respectively according to their binding signals, C-scores, and BC-scores. For each ranking, the top 250 sorted binding sites were used to infer binding motifs. For a TF with available binding sites less than 500, we used the top 50% of its sorted binding sites for motif inference. The genomic coordinates of the top binding sites were converted to DNA sequences using BEDTools. From a set of DNA sequences, we used DREME (v4.11.4) [18], MEME (v4.11.4) [23], and HOMER (v4.10.4) [41] to infer binding motifs. To make the inferred motifs comparable across the three software tools, we required each tool to report motifs with lengths of eight. In addition, we used the default setting for other parameters. For each TF, the top five inferred motifs from each tool were kept for evaluation. If any software could only identify less than five motifs, we took the smallest number of motifs across the three software tools.

The inference accuracy for a TF was first evaluated by the number of the five inferred motifs that were similar to the known motifs of the TF (*P* < 0.05; TOMTOM (v4.11.4) [55]). Using this measure, we evaluated the performance of different BC-score weights (*w*), binding site lengths (*l*), and their combinations for motif inference. The BC-score weight ranged from 0.1 to ten, with step lengths equal to 0.1 and 1 respectively for 0 to 1 and 1 to 10. The binding site length ranged from 20 to 400 with a step length equal to 20. For each sample (e.g., GM12878), to identify its optimal and robust parameters for motif inference, we divided its TFs with predicted binding sites and known motifs into five sets. With four sets as training data, we identified the optimal *w* and *l*. These parameters were evaluated on the other set of TFs as independent testing. We repeated this procedure by randomizing the five TF sets to estimate variance. In addition, we evaluated the motif inference of a TF using the number of its *in vitro* motifs that are significantly similar to the five inferred motifs of the TF.

### Motif entropy calculation

We calculated the entropy of a motif using its position weight matrix file (PWM). Each of the element in the matrix is denoted as *P*_*k,j*_, which is the frequency of the nucleotide *k* at the *j*th position of the motif. The *k* represents one of the four nucleotides. Therefore, the entropy of a motif is calculated as in Equation 1,

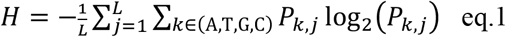

where *L* is the length of the motif.

### Gene expression and genome accessibility of GM12878 and K562 cell lines

The processed RNA-seq data and DNase-seq data for GM12878 and K562 cell lines were downloaded from the ENCODE portal (ENCSR000AEE, ENCSR109IQO, ENCSR000EMT, and ENCSR000EKQ). The genomic accessibility of a binding site is the average normalized read-depth signals within this binding site region, calculated by bigWigAverageOverBed (v2) from UCSC Kent tools.

## DATA AVAILABILITY

The code and data are available at https://github.com/gersteinlab/MOBI

## Supplementary figures

**Figure S1.**
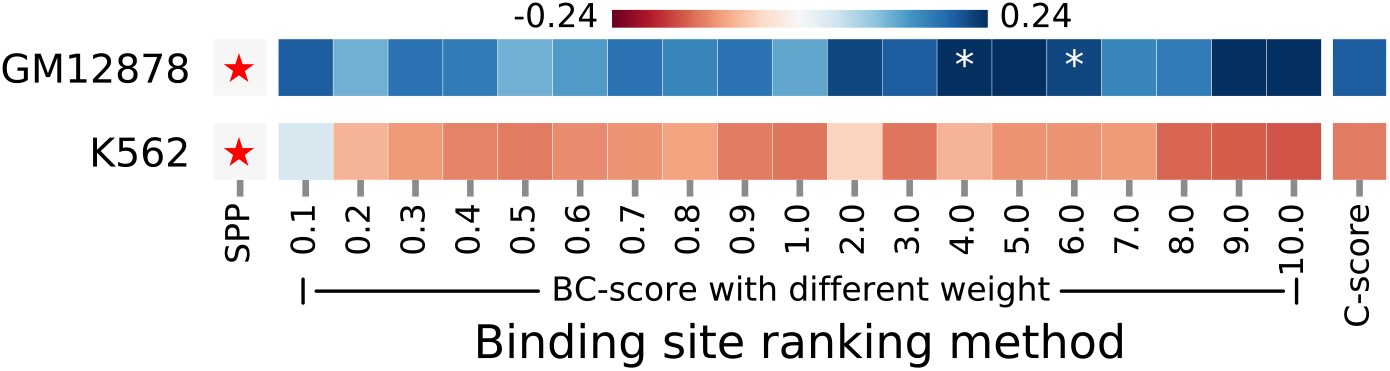
Transferability of C-scores across samples. The C-scores are calculated using binding sites detected from human cell lines and primary cells except GM12878 and K562. ‘*’ indicates statistical significance (*p* < 0.05) from Wilcoxon tests comparing the inference results from MOBI approaches to those from the conventional selections (red stars).

**Figure S2.**
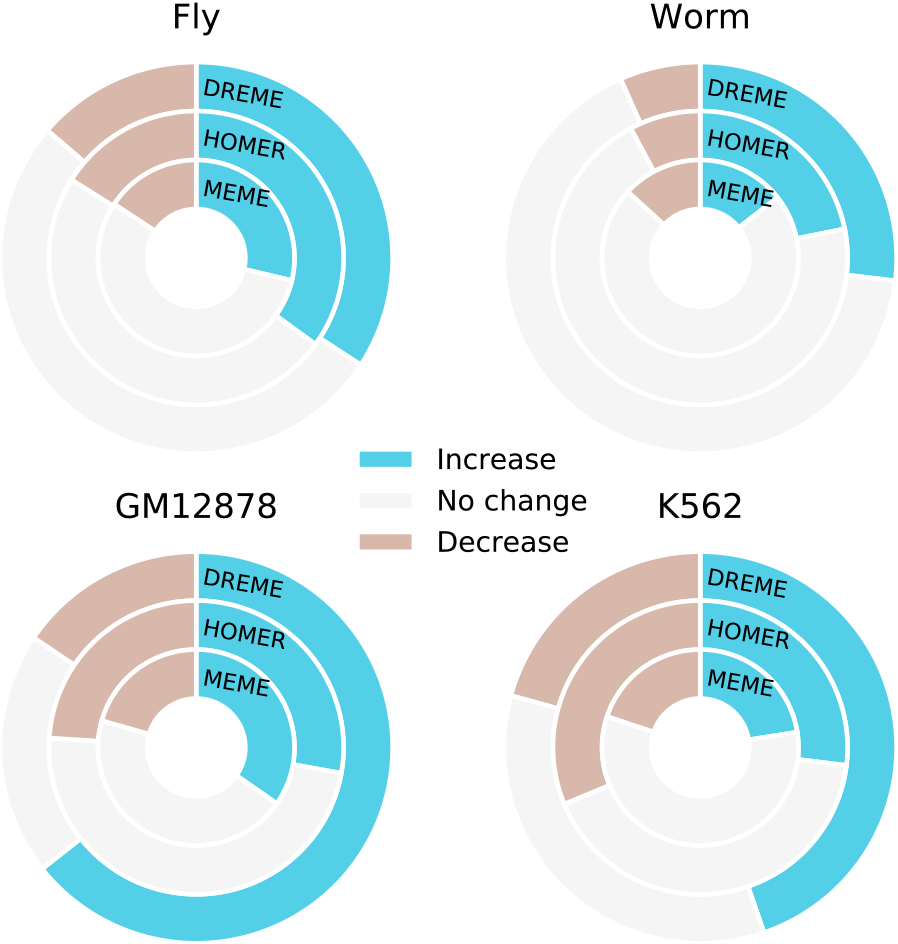
The fraction of TFs with improved motif inference using MOBI, compared to using conventional binding site selection.

**Figure S3.**
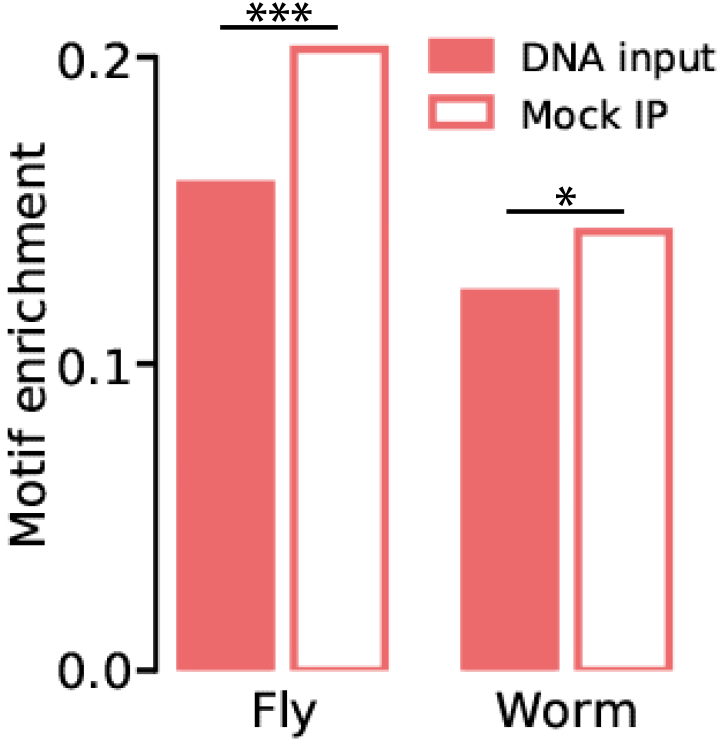
Compared to DNA input controls, mock IP controls detect binding sites (top 250 binding sites) enriched with more target motifs.

**Figure S4.**
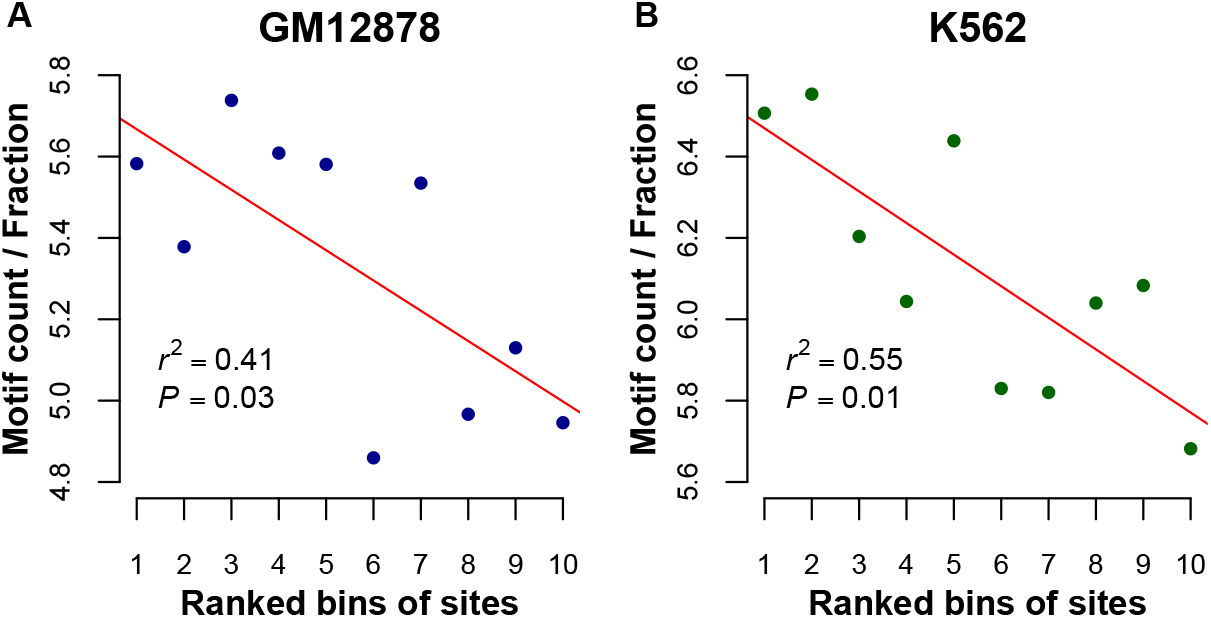
Association between the number of binding motifs used and the binding signal strength. The binding sites of a TF are ranked by their binding signals determined by ChIP-seq, in a descending order. Every fifty sites in the ranking are grouped into a bin. In each bin, the ratio between the motif count and the fraction indicates the number of motifs averaged over the sites with binding motifs. Each dot corresponds to the ratio of a bin averaged across the TFs. The red lines are linear regressions. The sample sizes of GM12878 (A) and K562 (B) are 52 and 87 TFs.

**Figure S5.**
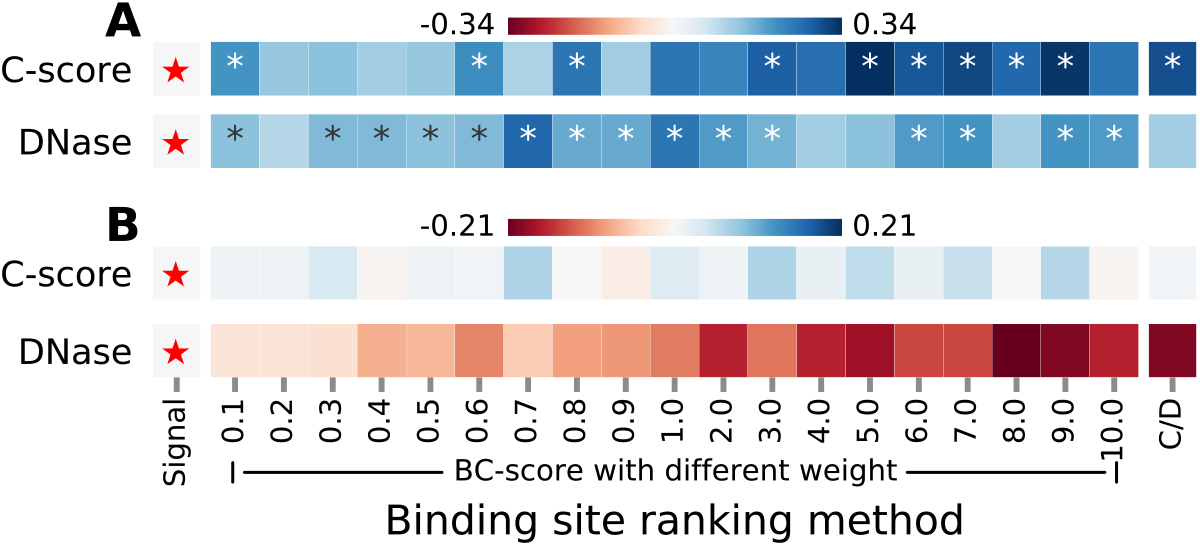
Comparing the effects of C-score and genomic accessibility on motif inference for TFs from GM12878 (A) and K562 (B). Genome accessibility are determined by DNase-seq respectively for the two cell lines. The accessibility of each binding site region is used as a proxy of C-score for motif inference. The red stars indicate the motif inference results from using the conventional binding site selection. These results are taken as the reference for the comparison to those from MOBI approaches respectively using C-scores and DNase accessibility. C/D respectively indicates using C-score or genome accessibility for binding site selection. ‘*’ indicates statistical significance (*p* < 0.05) from Wilcoxon tests.

## SURPPORTING INFORMATION

### Genome accessibility as a proxy of C-score for motif inference

We postulate that a genomic region with a large C-score likely has high accessibility, and thus the latter may be used as a proxy of the C-score. To this end, we focused on the simple samples, i.e. GM12878 and K562, which have DNase-seq data from the ENCODE portal. For each of the TF binding sites, its genomic accessibility is indicated by the number of reads mapped to it. Indeed, the genomic accessibility and C-scores of binding sites have modest correlations respectively for GM12878 (*ρ* = 0.682, *P* < 2.2 × 10^−16^) and K562 (*ρ* = 0.625, *P* < 2.2 × 10^−16^), indicating that highly accessible regions in the genome tend to have binding events of many different TFs. Note that these events observed by ChIP-seq may be spurious. Instead of the C-score, we used the genomic accessibility of a binding site for binding site selection, and then inferred motifs from the selected sites. For GM12878, a wide range of weights on genomic accessibility substantially improved motif inference, compared to conventional binding site selection (fig. S5A). However, for K562, since C-score alone already did not improve motif inference, using genomic accessibility as a proxy yielded no improvement (fig. S5B). For both of the cell lines, C-scores outperformed genomic accessibility for motif inference (fig. S5).

### Association between binding motif abundance and binding signal strength

We postulate that the strong binding signal of a site is potentially due to many binding motifs existing in the site. To test this, for each TF, we focused on the top 500 binding sites with strongest binding signals determined by ChIP-seq, and then grouped these ranked sites into 10 bins with the first bin containing the strongest binding sites. For each bin, we calculated the fraction of the sites having binding motifs (*p* < 10^−4^) and the number of binding motifs averaged over all the binding sites, herein referred to as motif count. The ratio between the motif count and the fraction corrects the bias that the bin of stronger binding sites has higher prevalence of binding sites with binding motifs. This ratio indicates the average number of binding motifs per site carrying binding motifs. However, this ratio is sensitive to the accuracy of the binding sites detected by ChIP-seq, therefore we focused on the top strongest 500 binding sites of TFs from GM12878 and K562. We observed strong linear relationship between the average ratio across the TFs and the bin ranks for both GM12878 (Fig. S4A) and K562 (Fig. S4B).

